# VCAP-102 Achieves Exceptional BBB Penetration in Marmosets After Ten Years of AAV Capsid Evaluation

**DOI:** 10.64898/2025.12.06.692617

**Authors:** Yasunori Matsuzaki, Ayumu Konno, Yasuo Uchida, Tetsuya Terasaki, Hirokazu Hirai

## Abstract

Efficient passage of adenoassociated virus (AAV) vectors across the blood–brain barrier (BBB) is essential for achieving broad central nervous system transduction following systemic delivery. Since PHP.B was reported in 2016 to exhibit high BBB permeability in C57BL/6J mice, many BBB-penetrant capsids have been developed, yet their performance often varies substantially across species.

From 2016 to 2025, we produced each newly reported BBB-penetrant capsid as its sequence became available and evaluated it under standardized intravenous administration conditions in marmosets. This included PHP.B, PHP.eB, AAV-F, CAP-B22, CAP-B10, CPP.16, CAP-Mac, and BR1N. Although several of these capsids have been reported to penetrate the marmoset BBB, none reproduced these effects under our unified evaluation conditions. We also pursued several directed-evolution approaches to engineer AAV9-based variants with improved BBB permeability; however, none surpassed AAV9.

In 2025, VCAP-102 was reported as an AAV9-derived capsid with BBB permeability in mice and non-human primates. When evaluated using our standardized framework, VCAP-102 consistently mediated markedly stronger and more widespread whole-brain transduction than AAV9 across all three animals tested.

These findings provide a decade-long, systematic assessment of BBB-penetrant AAV capsids in marmosets and identify VCAP-102 as a highly promising candidate for future systemically delivered AAV-based gene therapies.

## Introduction

In humans, the brain is the second heaviest organ after the liver and requires a continuous supply of oxygen and glucose delivered through an extensive vascular network. The blood– brain barrier (BBB) tightly regulates molecular exchange between the circulation and the brain parenchyma. Efficient gene delivery across the BBB is therefore essential for developing transformative therapies for neurological diseases with widespread pathology, such as Alzheimer’s disease and inherited neurodegenerative disorders.

Among naturally occurring serotypes, adeno-associated virus 9 (AAV9) exhibits relatively high blood–brain barrier (BBB) permeability. However, even high-dose intravenous administration of recombinant AAV9 results in <1% gene transduction in the mouse brain. In 2016, Gradinaru and colleagues developed PHP.B by inserting a seven–amino acid peptide into the AAV9 variable region VIII (VR-VIII) using the CREATE platform, demonstrating exceptional BBB penetration in C57BL/6J mice ^1^. Yet, our evaluation revealed that PHP.B did not improve brain transduction in marmosets ^2^—a result not previously reported, highlighting a pronounced species barrier. PHP.B was later shown to rely on Ly6A ^3,4^, a receptor not expressed in primates, thereby limiting its translational potential.

Following this, whenever a new BBB-penetrant capsid was described, we obtained its sequence upon publication, produced the corresponding vector, and evaluated it side-by-side in our standardized marmoset intravenous administration system. These variants—including CAP-B22, CAP-B10 ^5,6^, CPP.16 ^7^, and CAP-Mac ^8^—showed no advantage over AAV9 in our assays.

In 2024, Voyager Therapeutics identified 13 candidate VR-IV insertion sequences using the TRACER platform ^9^. Among them, VCAP-102, containing the HDSPHKSG insertion, was reported in 2025 to show robust BBB penetration in mice and non-human primates (NHPs) ^10^. When tested in our established marmoset system, VCAP-102 produced markedly stronger and more widespread whole-brain transduction than AAV9 in all three animals examined.

Here, we present a decade-long systematic evaluation of BBB-penetrant AAV capsids in marmosets and demonstrate that VCAP-102 is the first variant to achieve outstanding BBB permeability in this primate species.

## Results

### AAV capsid variants reported to efficiently cross the BBB in mice

BBB-penetrant AAV capsid variants reported since 2016 and subsequently evaluated in our marmoset system are summarized in Fig. 1A and Table S1. In 2016, Gradinaru et al. identified PHP.B—an AAV9 variant carrying a seven–amino acid insertion in VR-VIII—using the CREATE platform, demonstrating exceptional BBB permeability in C57BL/6J mice ^1^. We immediately tested PHP.B in marmosets by intravenously administering PHP.B- or AAV9-GFP, but observed no enhancement in brain transduction at 4 weeks ^2^.

**Figure 1.**
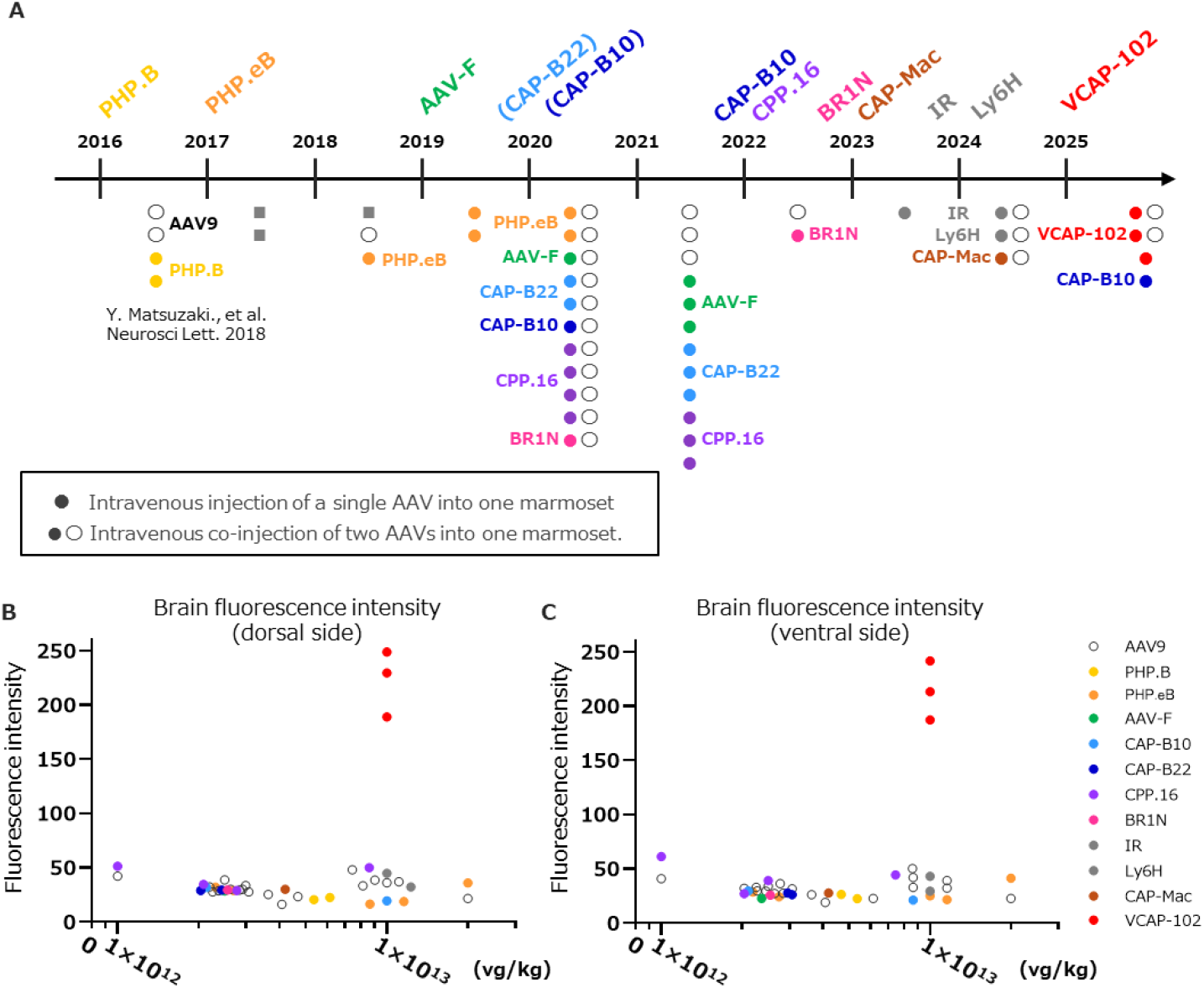
Systematic evaluation of BBB-penetrant AAV capsid variants in marmosets over the past decade. (A) Timeline summarizing BBB-penetrant AAV capsid variants reported between 2016 and 2025 and evaluated in our marmoset intravenous injection system. Each newly reported AAV capsid (e.g., PHP.B, PHP.eB, AAV-F, CAP-B22, CAP-B10, CPP.16, BR1N, CAP-Mac, VCAP-102) was produced upon publication and assessed under unified conditions. The gray squares in 2017–2018 indicate the marmosets used for our screening based on a modified CREATE method. The gray circles in 2023–2024 represent the marmosets that received intravenous administration of AAV9 capsid variants targeting the insulin receptor (IR) or Ly6H, which we identified as candidate endothelial receptor targets. Filled circles (•) represent intravenous injection of a single AAV vector into one marmoset, whereas half-filled circles (• ○) indicate co-injection of two AAV vectors into the same animal. (B, C) Brain-wide fluorescence intensity (0–255 a.u.) measured 4 weeks after intravenous AAV administration. (B) Dorsal view and (C) ventral view of whole brains from marmosets injected with each capsid variant. Fluorescence intensities for AAV9 and AAV-F (green), CAP-B10 (blue), CAP-B22 (cyan), CPP.16 (purple), BR1N (magenta), CAP-Mac (brown), and VCAP-102 (red) are plotted as a function of viral dose (vg/kg). Although most capsids exhibited fluorescence intensities comparable to AAV9, VCAP-102 consistently produced markedly higher brain-wide fluorescence, with values reaching the upper saturation limit (255 a.u.). This indicates exceptionally efficient BBB penetration of VCAP-102 relative to previously reported capsid variants.

We further showed that PHP.B fails to outperform AAV9 in several mouse strains, including BALB/c ^11^. Later studies revealed that PHP.B utilizes Ly6A as its endothelial receptor, and strains lacking endothelial Ly6A expression do not exhibit BBB permeability ^3,4^.

Subsequent capsids reported to cross the BBB in mice—including PHP.eB ^12^, AAV-F (which penetrates the BBB in both C57BL/6 and BALB/c) ^13^, and the AAV2-based variant BR1N ^14^— were also examined in our marmoset system. None exhibited BBB permeability greater than AAV9 (Fig. 1B, C and Table S2).

### AAV capsid variants reported to cross the BBB in nonhuman primates

Since 2020, several capsids reported to penetrate the BBB in NHPs—including CAP-B22, CAP-B10 ^5,6^, CPP.16 ^7^, and CAP-Mac ^8^—have been successively developed (Table S1). For each newly reported capsid, we rapidly produced the corresponding AAV vector and tested its efficacy following intravenous administration in marmosets (Fig. 1A). Whole-brain GFP fluorescence showed no notable improvement relative to AAV9 for any variant (Fig. 1B, C and Table S2).

We further quantified AAV genomic DNA (gDNA) in the cerebral cortex and cerebellum for AAV-F, CAP-B22, and CPP.16. gDNA levels were comparable to AAV9 for all variants, and cortical levels were significantly lower for AAV-F (Fig. S1).

### Independent engineering of AAV9 variants by our group

In addition to evaluating externally developed capsid variants, we also undertook multiple independent engineering efforts to improve the BBB permeability of AAV9 in marmosets.

First, in 2017–2018, we modified the CREATE method originally used to identify PHP.B ^1^ and adapted it for systemic screening in marmosets. We generated an AAV9 capsid library containing random seven–amino acid insertions in VR-VIII, administered the library intravenously, and two weeks later injected a mixture of neuron-specific and astrocyte-specific Cre-expressing AAVs ^15,16^ into the brain parenchyma (gray squares in Fig. 1A). This strategy was designed to recover recombined variants that had successfully crossed the BBB and reached neuronal and glial populations. However, no unique candidate BBB-penetrant variants were isolated.

Next, in 2023–2024, guided by proteomic analyses identifying insulin receptor (IR) and Ly6H as highly expressed on marmoset brain endothelial membranes ^17^, we established HEK293T cell lines stably expressing IR or Ly6H and screened for AAV9 variants with increased infectivity toward these membrane targets. Although two candidate variants were selected (AAV-IR and AAV-Ly6A), neither increased brain GFP expression following intravenous administration, indicating no detectable improvement in BBB permeability in marmosets (Fig. 1 and Table S2).

### VCAP-102 shows uniquely high BBB permeability in marmosets

In 2025, Nonnenmacher et al. reported VCAP-102, an AAV9-derived variant carrying the HDSPHKSG insertion in VR-IV, which efficiently crosses the BBB in both mice and NHPs ^10^. We generated VCAP-102 and assessed its performance in three marmosets. Four weeks after intravenous administration, whole-brain fluorescence imaging revealed exceptionally strong and widespread GFP expression—far exceeding that of AAV9 and all previously evaluated variants (Fig. 1 and Fig. 2).

**Figure 2.**
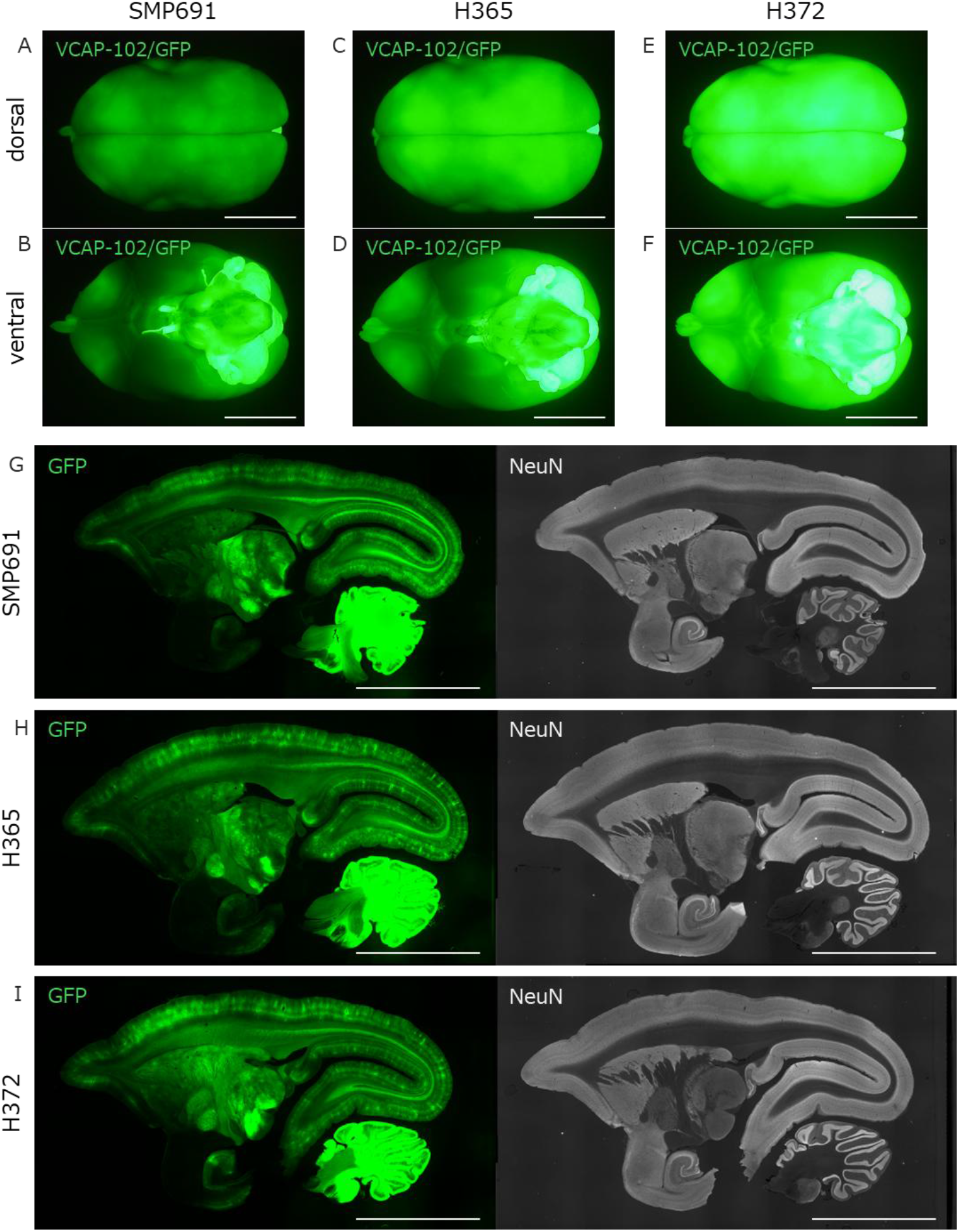
Whole-brain and sectional fluorescence images of marmosets intravenously injected with VCAP-102. (A–F) Whole-brain dorsal (A, C, E) and ventral (B, D, F) fluorescence images taken 4 weeks after intravenous injection of VCAP-102-GFP in three marmosets (SMP691, H365, H372). All animals showed extremely strong and widespread GFP fluorescence, markedly exceeding that observed with any other BBB-penetrant capsid variants tested in this study. Scale bars; 10 mm. (G–I) Sagittal brain sections from the same three animals showing robust GFP expression throughout cortical and subcortical regions. Corresponding NeuN-stained sections (right panels) are shown for anatomical reference. Scale bars; 10 mm. These findings corroborate the whole-brain observations reported in the main text and demonstrate that VCAP-102 achieves remarkably high levels of gene expression across the marmoset brain.

### BBB permeability assessed by co-administration with AAV9

To directly compare BBB permeability between each capsid variant and AAV9 within the same animal, we performed co-administration experiments in which the variant and AAV9 expressed different fluorescent proteins. For all variants except VCAP-102, the capsid variants expressed mCherry, while the control AAV9 expressed GFP (Fig. 3A-T).

**Figure 3.**
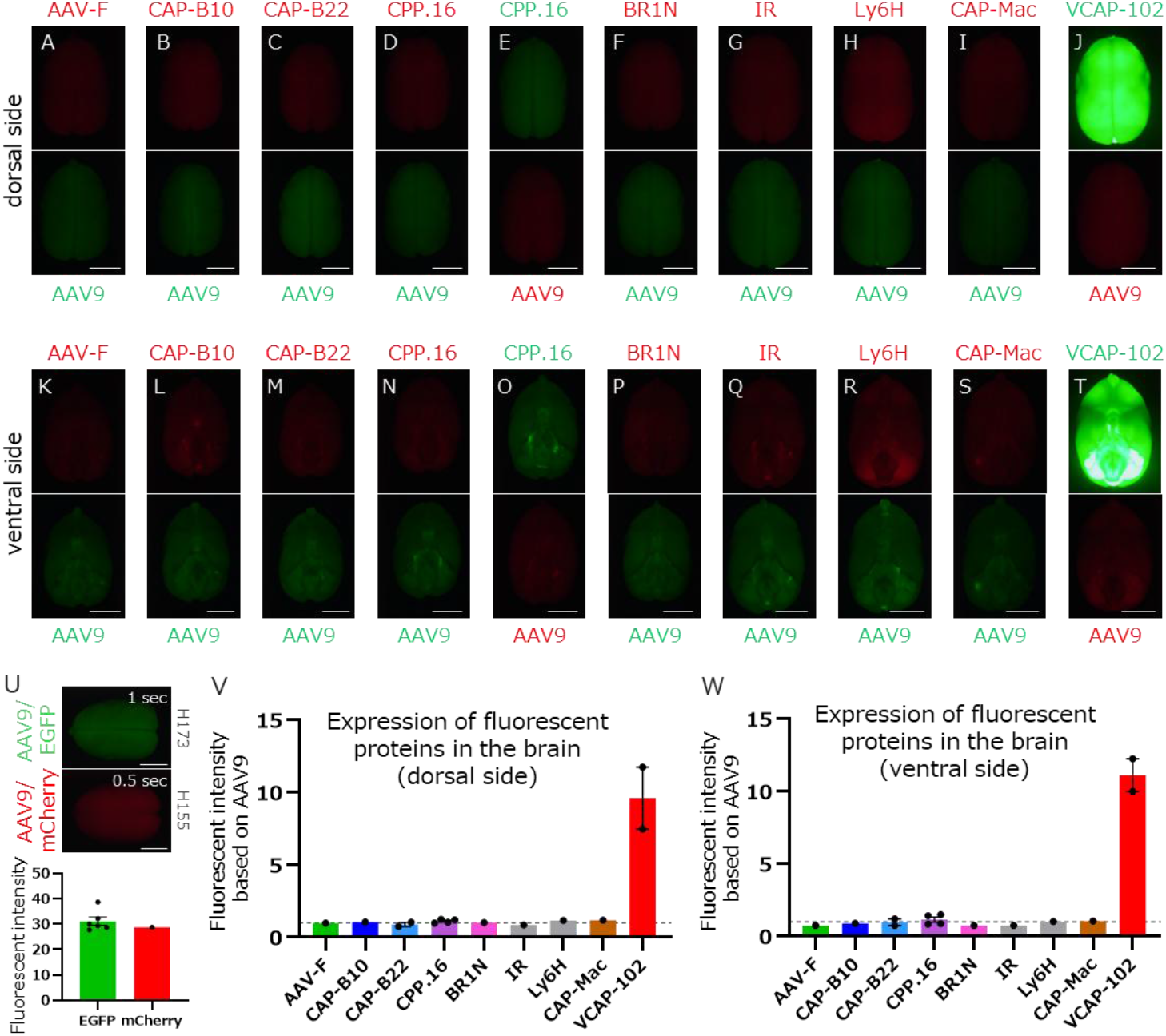
Co-injection assay comparing BBB permeability of AAV capsid variants with AAV9 in marmosets. (A–T) Representative dorsal (A–J) and ventral (K–T) whole-brain fluorescence images obtained 4 weeks after intravenous co-administration of each AAV capsid variant (expressing mCherry) together with AAV9 (expressing EGFP). For CPP.16, the reciprocal fluorophore combination (CPP.16–EGFP with AAV9–mCherry) was also evaluated (E, O). For VCAP-102, the fluorophore pairing was intentionally reversed (VCAP-102–EGFP with AAV9–mCherry) to allow accurate quantification of the markedly stronger fluorescence produced by VCAP-102 (J, T). Scale bars; 10 mm. (U) Control experiment showing that AAV9–EGFP (1-s exposure time) and AAV9–mCherry (0.5-s exposure time) produce comparable whole-brain fluorescence levels, validating the exposure settings used throughout the study. (V, W) Fluorescence intensity ratios of each variant relative to co-administered AAV9 in the same animal, calculated separately for the dorsal (V) and ventral (W) brain surfaces. All capsid variants except VCAP-102 exhibited fluorescence ratios near 1, indicating BBB permeability equivalent to AAV9. In contrast, VCAP-102 showed >10-fold higher fluorescence than AAV9 in both animals tested, confirming its uniquely strong BBB penetration in marmosets.

For VCAP-102, this fluorophore pairing was intentionally reversed—VCAP-102 expressed GFP and the control AAV9 expressed mCherry—to ensure accurate quantification of the markedly stronger fluorescence produced by VCAP-102 (i.e., AAV9-mCherry co-administered with VCAP-102-GFP) (Fig. 3J, T). For CPP.16, we additionally evaluated the reciprocal combination (AAV9-mCherry with CPP.16-GFP) (Fig. 3E, O).

When AAV9-GFP and AAV9-mCherry were administered under identical conditions, whole-brain fluorescence levels were comparable between GFP (1-s exposure) and mCherry (0.5-s exposure) (Fig. 3U and Fig. S2). Therefore, throughout all experiments, GFP and mCherry images were acquired at 1 s and 0.5 s exposure times, respectively.

Using the corresponding whole-brain dorsal (or ventral) fluorescence intensity from AAV9-injected animals as the reference, we calculated variant-to-AAV9 fluorescence ratios. All variants except VCAP-102 exhibited ratios near 1 (Fig. 3V, W), indicating BBB permeability equivalent to AAV9. In contrast, the two animals co-administered with AAV9 exhibited the VCAP-102/AAV9 fluorescence ratio greatly exceeded 10-fold (Rightmost bars in Fig. 3V, W).

Fluorescence values were recorded on a relative scale of 0–255. Importantly, the exposure times used in this study—1 s for GFP and 0.5 s for mCherry—represent the lowest settings at which both fluorophores remain simultaneously detectable. When either exposure time is further reduced, the fluorescence signals for both GFP and mCherry fall below the detection threshold, making these settings the practical lower limit for dual-channel imaging.

Even under these minimum-detection exposure conditions, brain regions transduced by VCAP-102 displayed clear GFP saturation (insets in Fig. S3B-F). Thus, reducing exposure time to avoid GFP saturation would completely eliminate detectable mCherry signal, preventing a valid comparison within the same animal. Accordingly, the true fluorescence intensity of VCAP-102 relative to AAV9 is likely substantially higher than the ratios shown in Fig. 3V and 3W.

## Discussion

In this study, we systematically evaluated the brain transduction efficiency of BBB-targeted AAV capsid variants in marmosets under standardized conditions for approximately ten years, beginning with the initial report of the BBB-penetrant variant PHP.B in 2016. Each time a new variant was reported, we generated the corresponding AAV vector expressing a fluorescent protein under the CBh promoter and assessed its ability to deliver genes to the marmoset brain following intravenous administration.

In addition to testing externally developed variants, we also pursued independent engineering strategies. In 2017–2018, we adapted the CREATE method for use in marmosets and performed in vivo screening in three animals; however, this approach did not yield unique variants capable of efficiently crossing the BBB. In 2023–2024, we conducted an independent directed-evolution screen that combined a VR-VIII insertion library with HEK293T cell lines stably expressing membrane proteins highly enriched in marmoset brain endothelial cells (IR and Ly6H). Sixteen candidate variants were selected and evaluated in marmosets, but none exhibited BBB permeability exceeding that of AAV9.

Through these multi-step efforts, we ultimately identified—within our marmoset model— VCAP-102 as the first variant to demonstrate markedly enhanced BBB penetration and robust whole-brain transduction in 2025.

Previous studies reported that CAP-B10, CPP.16, and CAP-Mac achieve approximately 5–12-fold higher gene delivery efficiency than AAV9 in the marmoset brain based on mRNA quantification ^6–8^. In contrast, VCAP-102 has been reported to reach exceptionally high levels, approximately 280–500-fold above AAV9 ^10^. The striking whole-brain GFP expression we observed with VCAP-102 in this study is fully consistent with these prior findings.

At the same time, although CAP-B10, CPP.16, and CAP-Mac have been reported to outperform AAV9 by 5–12-fold, our evaluations showed transduction efficiencies that were essentially comparable to AAV9. We believe it is premature to conclude that these variants inherently lack enhanced BBB permeability in marmosets, as several factors may underlie this discrepancy.

First, differences in AAV production and recovery methods may influence receptor binding and BBB transport. In this study, we used AAV particles naturally released into serum-free culture supernatant, whereas many prior studies relied on cell-lysis–based recovery methods. Because AAV capsids carry glycans and other post-translational modifications (PTMs) and the PTM landscape varies across manufacturing platforms ^18^, such methodological differences could alter capsid–receptor interactions. In addition, many earlier studies quantified transduction based on mRNA extracted from brain tissue, whereas our analysis relied primarily on whole-brain fluorescence, representing a fundamentally different readout.

Second, CPP.16 exhibits concentration-dependent aggregation ^7^, and we also observed subtle but clearly detectable aggregates. Even microscale aggregation can markedly impair BBB permeability, and similar aggregation-related effects cannot be excluded for the other variants.

Third, genetic background differences between marmoset colonies may contribute. As PHP.B shows dramatic strain-dependent differences between C57BL/6J and BALB/c mice ^3,4,11^, marmoset colonies maintained in Japan and those used in the United States differ genetically, which may influence receptor expression or BBB physiology. The colony used in this study has been maintained for 15 years with periodic introduction of ~30 new animals across six events, yet remains relatively homogeneous in kinship structure.

Taken together, it cannot be ruled out that the results of this study may have been influenced by both the AAV manufacturing workflow and the characteristics of the marmoset cohort investigated here. Therefore, even variants that exhibited BBB permeability comparable to AAV9 in our experiments may show higher transduction efficiency when produced using alternative AAV manufacturing protocols or when tested in genetically more diverse marmoset populations.

Finally, although previously reported BBB-penetrant AAV variants have shown inconsistent performance across studies and species, our data demonstrate that VCAP-102 uniquely achieves robust BBB penetration and widespread brain transduction in marmosets. Together with prior evidence identifying ALPL (tissue-nonspecific alkaline phosphatase) as its endothelial receptor—a molecule highly conserved from rodents to primates, including humans—these findings position VCAP-102 as one of the most promising candidates for future systemic gene delivery to the human brain.

## MATERIALS & METHODS

Please refer to supplemental materials and methods.

## DATA AVAILABILITY

All data supporting the findings of this study are available within the article and its supplemental files. Raw data can be provided upon request.

## ACKNOWLEDGMENTS

This work was supported by grants from the Program for Brain Mapping by Integrated Neurotechnologies for Disease Studies (Brain/MINDS; JP20dm0207057/JP21dm0207111 [to H.H.]) and Multidisciplinary Frontier Brain and Neuroscience Discoveries (Brain/MINDS 2.0; JP24wm0625103 [to H.H.]) from the Japan Agency for Medical Research and Development (AMED); and by MEXT/JSPS KAKENHI (24K15743 [to Y.M.], 22K06454/24H01221 [to A.K.], and 23H02791 [to H.H.]). The authors thank Asako Ohnishi, Nobue McCullough, Chieko Miyazawa, Ayako Sugimoto, and Keiko Sato for AAV vector production; Motoko Uchiyama, Minako Noguchi, and Yoshiko Nomura for marmoset care and management; Junko Sugi for immunohistochemistry; and Minako Noguchi also for creating the marmoset illustration used in the graphical abstract.

## AUTHOR CONTRIBUTIONS

All authors contributed to the overall experimental design.

Y.M. performed the intravenous AAV injections in marmosets, carried out immunohistochemistry, and conducted data analysis.

A.K. generated the AAV vectors and conducted all screening experiments using the AAV capsid variant libraries in both marmosets and cultured cells.

Y.U. and T.T. proposed candidate membrane proteins enriched in marmoset brain vascular endothelial cells based on their own research findings and helped guide the conceptual direction of the study.

H.H. supervised the project, provided oversight for data interpretation, edited the manuscript, and finalized the study.

## SUPPLEMENTAL INFORMATION

Figures S1–S3 and Tables S1–S2

