## Supplemental figure S1-S3 for "VCAP-102 Achieves Exceptional BBB Penetration in Marmosets After Ten Years of AAV Capsid Evaluation"

**Supplemental figures**

**Figure S1.**

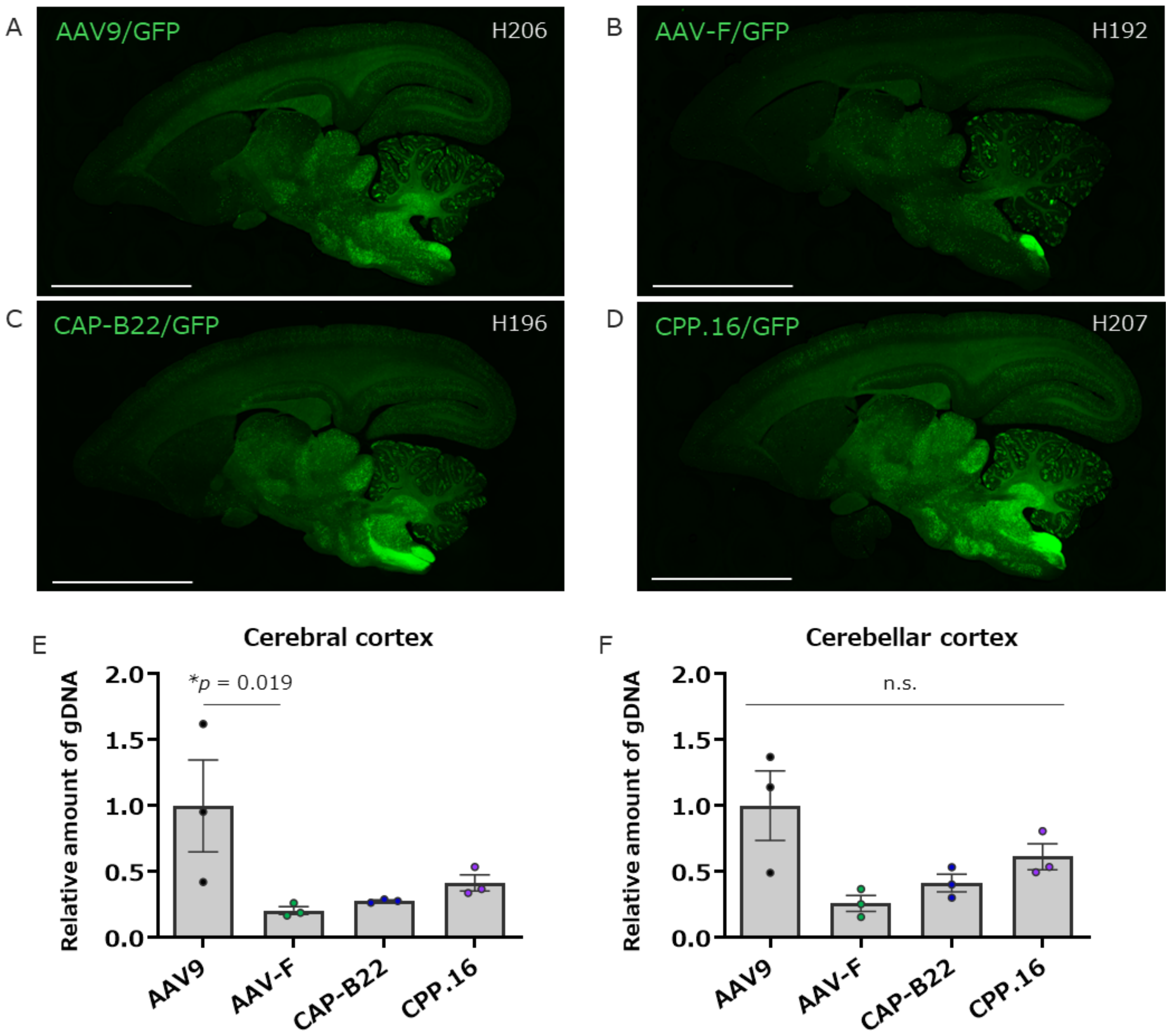

**Figure S1. Quantification of AAV genome DNA levels in the marmoset brain following intravenous injection of AAV-F, CAP-B22, or CPP.16.**

(A–D) Representative coronal brain images from marmosets injected with AAV9-GFP (A), AAV-F-GFP (B), CAP-B22-GFP (C), or CPP.16-GFP (D). Scale bars; 10 mm.

(E, F) Quantification of AAV genomic DNA (gDNA) in the cerebral cortex (E) and cerebellar cortex (F) by qPCR. gDNA levels for each capsid variant were normalized to those of AAV9. Consistent with the results described in the main text, none of the three variants (AAV-F, CAP-B22, CPP.16) exceeded AAV9 in cortical gDNA levels, and AAV-F showed significantly lower levels than AAV9 in the cerebral cortex ($p = 0.019$). No significant differences were observed among capsids in the cerebellar cortex. Error bars indicate the SEM, and each dot represents the value obtained from an individual marmoset.

**Figure S2.**

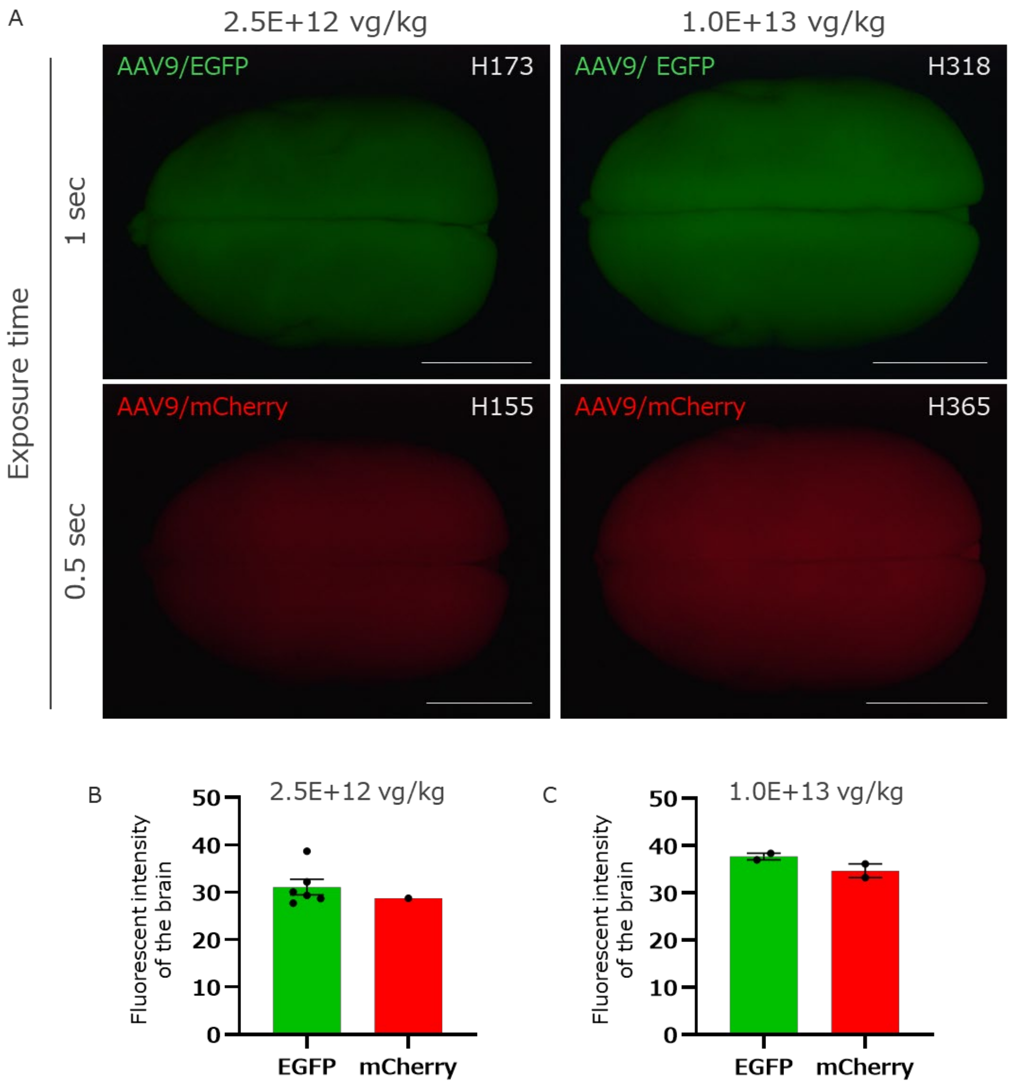
A
2.5E+12 vg/kg
1.0E+13 vg/kg
Exposure time
1 sec
0.5 sec
AAV9/EGFP
H173
AAV9/ EGFP
H318
AAV9/mCherry
H155
AAV9/mCherry
H365
B
2.5E+12 vg/kg
Fluorescent intensity of the brain
50
40
30
20
10
0
EGFP
mCherry
C
1.0E+13 vg/kg
Fluorescent intensity of the brain
50
40
30
20
10
0
EGFP
mCherry

**Figure S2. Comparison of whole-brain fluorescence intensity between AAV9-GFP and AAV9-mCherry at different exposure times.**

(A) Whole-brain fluorescence images of marmosets intravenously injected with AAV9-GFP (1 s exposure) or AAV9-mCherry (0.5 s exposure) at doses of $2.5\times10^{12}$ vg/kg (left) or $1.0\times10^{13}$ vg/kg (right). Scale bars; 10 mm.

(B, C) Quantification of brain fluorescence intensity at each dose. AAV9-GFP (1 s exposure) and AAV9-mCherry (0.5 s exposure) showed comparable whole-brain fluorescence levels. Therefore, GFP and mCherry signals were imaged using 1 s and 0.5 s exposure times, respectively, throughout this study.

**Figure S3.**

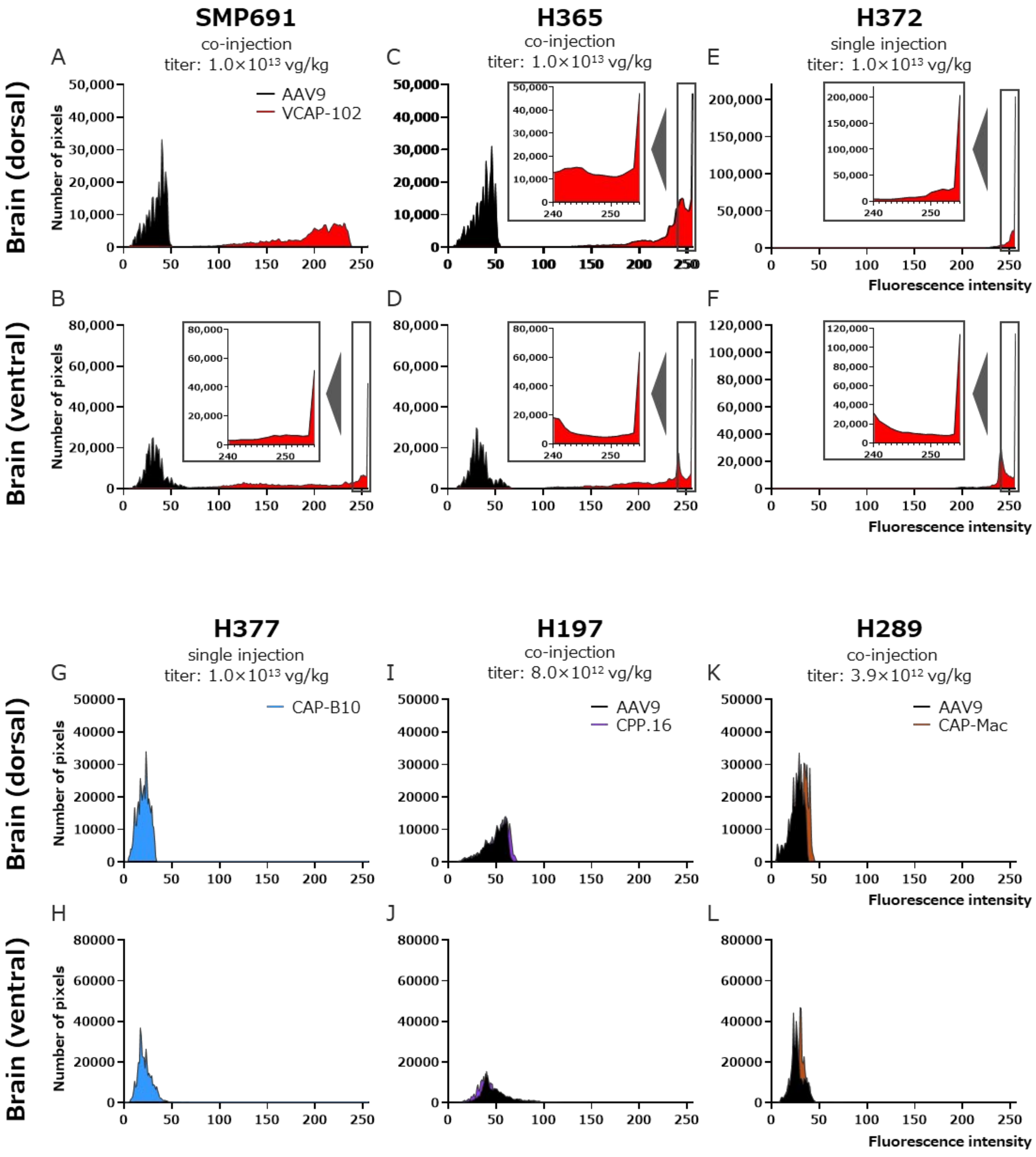
SMP691
co-injection
titer: 1.0×10¹³ vg/kg
H365
co-injection
titer: 1.0×10¹³ vg/kg
H372
single injection
titer: 1.0×10¹³ vg/kg
A
B
C
D
E
F
Brain (dorsal)
Brain (ventral)
Number of pixels
AAV9
VCAP-102
Fluorescence intensity
H377
single injection
titer: 1.0×10¹³ vg/kg
H197
co-injection
titer: 8.0×10¹² vg/kg
H289
co-injection
titer: 3.9×10¹² vg/kg
G
H
I
J
K
L
CAP-B10
AAV9
CPP.16
AAV9
CAP-Mac

**Figure S3. Whole-brain fluorescence intensity histograms of AAV-injected marmosets.**

Whole-brain dorsal (A, C, E, G, I, K) and ventral (B, D, F, H, J, L) fluorescence intensity histograms obtained three to four weeks after intravenous AAV administration in six marmosets (SMP691, H289, H365, H372, H377, H197). AAV9, VCAP-102, CAP-B10, CPP.16, and CAP-Mac were evaluated either by single injection or by co-injection, as indicated for each animal. Fluorescence intensity was recorded as relative values ranging from 0 to 255. For capsids other than VCAP-102, exposure times were minimized to the detection threshold (GFP: 1 s, mCherry: 0.5 s).

Insets in panels B–F show enlarged views of the upper-intensity range (240–255), highlighting the distribution of pixels near the saturation level. Because saturated pixels were recorded as 255, all brain regions in which VCAP-102 expression exceeded the dynamic range of detection appear as a vertical accumulation at this value.

Despite using shortened exposure conditions, all three marmosets injected with VCAP-102 exhibited strong GFP expression with signals saturating the upper limit (255 a.u.) across multiple brain regions. As saturated values were stored as 255 in the histogram analysis, the actual fluorescence intensity of VCAP-102 relative to AAV9 is likely higher than the ratios shown in Figure 3.
