## Supplemental materials and methods for "VCAP-102 Achieves Exceptional BBB Penetration in Marmosets After Ten Years of AAV Capsid Evaluation"

### Animals

The present study included 44 common marmosets (*Callithrix jacchus*). All animals were bred and maintained at the Gunma University Bioresource Center. The marmosets were housed in breeding rooms under controlled environmental conditions, including temperature (27–30°C), humidity (25–45%), and a 12-h light/dark cycle. Filtered water was provided ad libitum.

The animals were fed 45–50 g of soaked monkey chow (CMS-1; CLEA Japan, Tokyo, Japan) supplemented with fruits, vegetables, soybean flour, quail eggs, or boiled chicken around noon. In addition, "marmoset dumplings" were provided on weekday afternoons around 3:00 p.m. These were prepared by mixing CMS-1 soaked in hot water with honey, oligosaccharides, milk powder, vitamin supplements, Lactobacillus powder, and gum arabic powder.

All housing and husbandry conditions complied with the Guide for the Care and Use of Laboratory Animals, 8th edition. Every effort was made to minimize animal suffering and to reduce the number of animals used. All procedures for animal care and experimentation were approved by the Japan Neuroscience Society ("Guidelines for Experiments on Primates in the Field of Neuroscience") and the Institutional Animal Care and Use Committee of Gunma University (approval Nos. 18-019, 20-053, 21-063, 23-018, 23-057, and 24-057).

### Construction of plasmids

The expression plasmids pAAV-CBh-EGFP-WPRE-hBGpA and pAAV-CBh-mCherry-WPRE-hBGpA were used to drive constitutive expression of EGFP or mCherry under the control of the CBh promoter [1]. The Kozak sequence and fluorescent protein genes were inserted immediately downstream of the CBh promoter into the pAAV expression plasmid at the AgeI and NotI restriction sites.

The AAV2/9 packaging plasmid (pAAV2/9), which contains the AAV2 rep gene and the AAV9 cap gene, was provided by James M. Wilson. The cap genes of PHP.B, PHP.eB, AAV-F, CAP-B10, CAP-B22, and VCAP-102 were generated based on AAV9. Humanized DNA sequences corresponding to the amino acid sequences reported in the original publications were incorporated into synthetic oligonucleotides and inserted into the appropriate sites of the respective cap genes in pAAV2/9. The AAV-CPP.16 plasmid was kindly provided by Dr. Fengfeng Bei (Harvard Medical School). The plasmid for producing CAP-Mac, pUCmini-iCAP-AAV.CAP-Mac, was kindly provided by Dr. Viviana Gradinaru (California Institute of Technology). The BR1 and BR1N plasmids, both derived from AAV2, were constructed using a similar strategy: the humanized insert sequence was synthesized as an oligonucleotide and subsequently introduced into the target site of pRC2-mi342 [2].

All gene engineering procedures were approved by the Institutional Committee of Gunma University (approval Nos. 15-038, 20-018, 23-056, 254-054, and 24-070).

**CREATE-based Screening of AAV Capsids**

An AAV capsid library was generated using the CREATE method as previously described [3]. The plasmids rAAV-Cap-in-cis-lox genome, pRep-AAP, and pRep-AAP deltaX/A used for generating the AAV library for CREATE screening were kindly provided by Drs. Viviana Gradinaru and Benjamin Deverman. Unlike in rodents, genetically engineered marmosets that express Cre in specific brain cell types are not readily available. Therefore, Cre expression was achieved using AAV vectors. Specifically, the AAV library was first administered intravenously through the femoral vein. Two weeks after the AAV library injection, a mixture of AAV9/NSE-GFP-P2A-Cre and AAV9/GFAP-GFP-P2A-Cre (mixed at a 1:1 ratio) was injected directly into the cerebral cortex. This order of administration was chosen to avoid the production of anti-AAV9 neutralizing antibodies (NAbs) prior to library delivery. If AAV9 vectors were injected into the brain first, rapid induction of NAbs against AAV9 could be elicited [4], potentially resulting in capture of all AAV9-based library particles by the NAbs upon subsequent intravenous administration. Three or four weeks after intraparenchymal injection of the Cre-expressing AAV vectors, GFP-positive cortical tissue was dissected under a fluorescent stereomicroscope. AAV genomes were extracted from the collected tissue, and PCR amplification was performed using primers designed to detect Cre-dependent inversion events.

**Screening for AAV capsids targeting membrane proteins highly expressed in marmoset endothelial cells**

To generate stable cell lines expressing the target membrane proteins, the common marmoset (*Callithrix* *jacchus*) insulin receptor (cjIR) and lymphocyte antigen 6H (cjLy6H) were cloned into the lentiviral expression vector pCL20c/CBh.*-P2A-GFP.WPRE using PCR and the In-Fusion HD Cloning Kit (TaKaRa Bio, Shiga, Japan). The following primer sets were used:

cjIR-F, 5′-TTCAGGTTGGACCGGTGCCACCATGGGAGCCGGGAGCCGCCGG-3′;

P2A-cjIR-R, 5′-CGTGGCTCCGGAGCCGGAAGGGTTGGAACGAGGCAAGGCC-3′;

cjLy6H-F, 5′-TTCAGGTTGGACCGGTGCCACCATGCTGCCTGCAGCCATGAAGGG-3′;

P2A-cjLy6H-R, 5′-CGTGGCTCCGGAGCCGAGCCCCGCCCAGAGGAGGG-3′.

VSV-G–pseudotyped lentiviral vectors were produced in HEK293T cells as previously described [5,6] and used to establish HEK293T cell lines stably expressing cjIR or cjLy6H on the plasma membrane.

Synthetic oligonucleotides containing 8 × NNK, corresponding to random eight–amino acid insertions (GenScript, Piscataway, NJ, USA), were inserted into the AAV9 capsid gene to generate a diversified AAV library, which was produced in HEK293T cells. The resulting AAV library was screened using the cjIR- or cjLy6H-expressing stable cell lines. For each target membrane protein, three rounds of screening were performed, and candidate insertion sequences presumed to enhance infection specificity were identified by next-generation sequencing (NGS) analysis. Each candidate was designated AAV9-IR or AAV9-Ly6H, respectively, and their blood–brain barrier permeability was subsequently examined in marmosets.

**Production of AAV vectors**

AAV vectors were collected from the culture supernatant. Recombinant single-stranded AAV vectors were produced in HEK293T cells (HCL4517; Thermo Fisher Scientific, Waltham, MA) using the previously described ultracentrifugation method [7]. Briefly, HEK293T cells were cultured in Dulbecco's Modified Eagle Medium (DMEM; D5796-500ML, Merck, Darmstadt, Germany) supplemented with 8% fetal bovine serum (26140-079, Sigma-Aldrich) at 37°C in 5% $CO_2$. Cells were transfected with three plasmids—the pAAV expression plasmid, pHelper (Stratagene, La Jolla, CA), and a rep/cap plasmid—using polyethylenimine "Max" (24765-1; Polysciences, Warrington, PA, USA).

Viral particles were harvested from the culture medium 6 days after transfection and concentrated by precipitation with 8% polyethylene glycol 8000 (PEG8000; Merck) and 500 mM sodium chloride. The precipitated vectors were resuspended in D-PBS(–) and purified by iodixanol density gradient ultracentrifugation (OptiPrep; Serumwerk Bernburg AG, Bernburg, Germany). The purified viral solution was then concentrated in D-PBS(–) using a Vivaspin 20 centrifugal concentrator (100,000 MWCO PES; Sartorius, Göttingen, Germany).

Genomic titers were determined by real-time quantitative PCR using a Thermal Cycler Dice Real Time System II TP900 or III TP970 (Takara Bio, Shiga, Japan) and Power SYBR Green PCR Master Mix (Thermo Fisher Scientific). The following primers targeting the WPRE sequence were used:
5′-CTGTTGGGCACTGACAATTC-3′ and 5′-GAAGGGACGTAGCAGAAGGA-3′.
The expression plasmid was used to generate a standard curve for absolute quantification.

AAV preparations were stored at 4°C for short-term use (up to a few months) and at −80°C for long-term storage.

**HEK293-T cell-based AAV** NAbs **assay**

A HEK293T cell–based NAbs assay for AAV2 or AAV9 was performed to select marmosets suitable for intravenous AAV administration. Whole blood was collected from candidate animals, and serum was isolated by centrifugation at 5,000 rpm for 5 min at 4°C. Serum samples were stored at 4°C for up to one week or at –80°C for long-term storage.

HEK293T cells were seeded at 4 × $10^4$ cells/well in 100 µL of DMEM supplemented with 10% FBS in a 96-well plate (μ-Plate 96 Well Square ibiTreat Sterilized; ibidi, Gräfelfing, Germany) and incubated for 2–6 h at 37°C in 5% $CO_2$.

AAV2/CBh-EGFP-WPRE-hBGpA or AAV9/CBh-EGFP-WPRE-hBGpA were diluted in serum-free DMEM to achieve optimized multiplicities of infection (MOIs) of $1 \times 10^3$ (AAV2) or $5 \times 10^4$ (AAV9). Marmoset serum was diluted **1:5** in serum-free DMEM. Equal volumes of diluted serum and titer-adjusted AAV2 or AAV9 were mixed in a 96-well plate (Nunc MicroWell 96-Well; Thermo Fisher Scientific) and incubated for 1 h at 37°C in 5% $CO_2$. The AAV/serum mixtures were then added to the HEK293T cells and incubated for 72–76 h under the same conditions.

EGFP fluorescence in HEK293T cells was imaged and quantified using the Image Cytometer Module of the BZ-X800 fluorescence microscope (Keyence, Osaka, Japan).

To assess NAbs activity, marmoset serum was pre-incubated with AAV2 or AAV9 mutant capsids as positive controls. As negative controls, AAV vectors known not to react with the serum were used. The absence of NAbs was defined as EGFP fluorescence ≥ 50% of the signal in the negative control, which served as the cut-off threshold.

**Intravenous injection of AAV vectors**

Each AAV vector was administered intravenously into the femoral vein of marmosets. Intravenous injections were performed using one of two methods: (1) connecting a winged needle and syringe via a three-way stopcock, or (2) directly attaching the syringe to the winged needle, as described previously [8]. After immobilization by intramuscular injection of a mixture of ketamine hydrochloride and xylazine hydrochloride, the AAV vector solution was immediately administered intravenously.

**Necropsy and whole-brain fluorescence imaging**

Marmosets were anesthetized with a cocktail of ketamine hydrochloride, xylazine hydrochloride, and isoflurane four to five weeks after viral injection (4.6 ± 0.2 weeks, mean ± S.E.M., n = 32). Animals were perfused transcardially with 300 mL of cold 1× PBS(–) containing 20 mM EDTA (311-90075, Nippon Gene, Tokyo, Japan), followed by 250 mL of cold 4% paraformaldehyde (PFA) in 1× phosphate buffer (PB). Brains were then removed.

Following necropsy, EGFP and mCherry fluorescence in whole marmoset brains was imaged using a Keyence VB-7010 fluorescence microscope. Fluorescence intensity analysis was performed using Fiji (ImageJ) [9].

**Immunohistology**

To examine EGFP expression in marmoset brain tissue, 100-µm-thick microtome sections were prepared and subjected to fluorescent immunostaining. Brains were bisected medially with a scalpel, the temporal lobes were trimmed, and the remaining tissue was embedded in 2% agarose gel. Sagittal sections (100 µm thick) were obtained using a microtome (VT1200S; Leica Microsystems GmbH, Wetzlar, Germany) and stored at 4°C in 1× PBS(–) containing $NaN_3$ until use.

Sections were stained using triple-fluorescence labeling, including nuclear staining with NucBlue (Hoechst 33342; Thermo Fisher Scientific). Tissue sections were incubated overnight at room temperature with the following primary antibodies diluted in blocking solution (2% donkey serum [S30-100ML; Merck, Darmstadt, Germany], BSA [A9647; Merck], 0.5% Triton X-100, and 0.03% $NaN_3$ in 1× PB):

- Rat monoclonal anti-GFP antibody (1:1,000; 04404-84; Nacalai Tesque, Kyoto, Japan)
- Mouse monoclonal anti-NeuN antibody (1:1,000; MAB377; Merck)

For visualization, sections were incubated for 3–4 **h** at room temperature in blocking solution containing the following secondary antibodies:

- Donkey anti-rat IgG Alexa Fluor Plus 488 (1:2,000; Thermo Fisher Scientific)
- Donkey anti-mouse IgG Alexa Fluor Plus 555 (1:2,000; Thermo Fisher Scientific)

Following secondary antibody incubation, sections were mounted on glass slides using ProLong Glass Antifade Mountant with NucBlue Stain (Thermo Fisher Scientific), allowed to cure, and stored at 4°C.

**Comparative quantification of viral DNA in the marmoset brains**

To quantitatively assess delivery of BBB-permeable AAV vectors to the marmoset brain, the amount of AAV genomic DNA was measured by qPCR. Marmosets were anesthetized with a cocktail of ketamine hydrochloride, xylazine hydrochloride, and isoflurane six weeks after viral injection (6.6 ± 0.1 weeks, mean ± S.E.M., n = 12). Marmosets were perfused with cold D-PBS(–) containing 20 mM EDTA, followed by 1% PFA in 1× PB. The left hemisphere was used for histological analysis, whereas the right hemisphere was used for AAV genomic DNA quantification. Following perfusion fixation, brains were rapidly removed. The left hemisphere was post-fixed in 4% PFA overnight at 4°C and sectioned into 100-μm-thick slices for immunostaining. Slices were stained using the rat monoclonal anti-GFP antibody (Nacalai Tesque) and Donkey anti-rat IgG Alexa Fluor Plus 488 (Thermo Fisher Scientific).

For viral DNA quantification, 20–35 mg of tissue was dissected from the cerebral cortex and cerebellar cortex of the right hemisphere. Tissue pieces were immersed in 30% ethanol on ice for at least 10 min, followed by immersion in 40% ethanol on ice for at least 10 min. PFA–ethanol displacement was performed by sequential immersion in ethanol solutions of increasing concentration (10% increments), ending with 99.5% ethanol. Ethanol was then completely evaporated by incubation at 50°C for 1 hour.

Genomic DNA was subsequently extracted using the Wizard Genomic DNA Purification Kit (Promega, Madison, WI). qPCR quantification was performed using the KAPA SYBR Fast qPCR Kit (Roche, Basel, Switzerland) on the Thermal Cycler Dice Real Time System II TP900 (Takara Bio). Primers were designed to span the EGFP and WPRE regions, using the following sequences:

- **qPCR-EGFP-For:** 5′-GGACGAGCTGTACAAGTAAAG-3′
- **qPCR-WPRE-Rev:** 5′-GGGAAGCAATAGCATGATACAAAGG-3′

**Statistical analysis**

Statistical analyses and graph generation were performed using GraphPad Prism versions 6 and 10 (GraphPad

Software, San Diego, CA). Data comprising multiple groups are presented as scatter plots (Figure 1). Bars indicate mean values, and error bars represent the standard error of the mean (SEM) (Figure 3; Figures S1 and S2). Each dot corresponds to the fluorescence measurement from an individual marmoset brain. Comparisons among multiple groups were performed using the Kruskal–Wallis test followed by Dunn's multiple-comparison test (Figure S1).
