## Supplemental table S1-S2 for "VCAP-102 Achieves Exceptional BBB Penetration in Marmosets After Ten Years of AAV Capsid Evaluation"

**Table S1. Summary of wild-type AAV serotypes and BBB-penetrant AAV capsid variants (2016–2025).**

| Name of AAV vector | AA452–458 (VR-IV) | AA587–590 (VR-VIII) | Proposal receptor | Method | Publication year |
|---|---|---|---|---|---|
| AAV9 (AAV hu.14) | NGSGQNQ | AQAQ | - | - | 2004 |
| AAV-PHP.B | NGSGQNQ | AQTLAVPFKAQ | Ly6A (mouse only) | CREATE | 2016 |
| AAV-PHP.eB | NGSGQNQ | DGTLAVPFKAQ | Ly6A (mouse only) | CREATE | 2017 |
| AAV-F | NGSGQNQ | AQFVVGQSYAQ | Ly6C1 (mouse only) | iTransduce | 2019 |
| AAV.CAP-B10 | DGAATKN | DGTLAVPFKAQ | (unknown) | M-CREATE | 2022 |
| AAV.CAP-B22 | DGQSSKS | DGTLAVPFKAQ | (unknown) | M-CREATE | 2022 |
| AAV.CPP.16 | NGSGQNQ | AQTVSALKAQ | (transcytosis by the cpp) | CPPs-insertion | 2022 |
| AAV.CAP-Mac | NGSGQNQ | AQLNTTKPIAQ | (unknown) | *in vivo* directed evolution | 2023 |
| VCAP-102 | NGHDSPHKSGQNQ | AQAQ | ALPL/TNAP | TRACER | 2025 |
| AAV2 | SGTTTQS | NRQA | - | - | 1996 |
| AAV-BR1 | SGTTTQS | QRGNRGTEWDAQA | (unknown) | random AAV display peptide libraries | 2016 |
| AAV-BR1N | SGTTTQS | NRGNRGTEWDAQA | (unknown) | BR1 variant (587N) | 2023 |

This table lists the amino acid sequences inserted into variable region IV (VR-IV; aa452–458 in AAV9) and variable region VIII (VR-VIII; aa587–590 in AAV9), along with proposed or validated endothelial receptors, engineering strategies, and publication years for each capsid. AAV9 (published in 2004) and AAV2 (1996) represent wild-type serotypes. BBB-penetrant AAV capsid variants have been reported from 2016 to 2025. BR1 is not a BBB-penetrant variant but an AAV2-based capsid that efficiently transduces brain endothelial cells; replacement of residue 587 (Q→N) generates BR1N, which acquires BBB permeability in mice.

Amino acids 452–458 are located within the VR-IV region, and amino acids 587–590 are located within the VR-VIII region. Receptor information is provided only when experimentally demonstrated or explicitly proposed in the original studies. “Method” indicates the engineering strategy used (e.g., CREATE, M-CREATE, TRACER, CPP insertion, or directed evolution). Among the listed variants, VCAP-102 is the only capsid with a confirmed endothelial receptor (ALPL/TNAP) in non-human primates.

### Table S2

**Table S2. Individual marmoset information and whole-brain dorsal and ventral fluorescence intensities following intravenous AAV administration.**

| Titer (vg/kg) | Individual ID | Name | AAV capsid | Fluorescent protein | Mean fluorescence intensity dorsal/ventral (a.u., 0–255) |
|---|---|---|---|---|---|
| $1.0 \times 10^{12}$ | H193 | Ginga | AAV9 | GFP | 42.2 / 40.8 |
| | | | CPP.16 | mCherry | 51.1 / 61.1 |
| $2.5 \times 10^{12}$ | H165 | Nogi | AAV9 | GFP | 27.5 / 29.4 |
| | | | PHP.eB | mCherry | 28.8 / 28.4 |
| | H184 | Tanpopo | AAV9 | GFP | 33.5 / 27.5 |
| | | | PHP.eB | mCherry | 32.0 / 24.2 |
| | H173 | Waboku | AAV9 | GFP | 28.6 / 29.7 |
| | | | AAV-F | mCherry | 28.4 / 22.6 |
| | H186 | Fuuwa | AAV9 | GFP | 30.1 / 32.8 |
| | | | CAP-B10 | mCherry | 31.8 / 29.5 |
| | H187 | Kaoru | AAV9 | GFP | 38.6 / 36.3 |
| | | | CAP-B22 | mCherry | 29.4 / 27.6 |
| | H183 | Uguisu | AAV9 | GFP | 30.1 / 31.6 |
| | | | CAP-B22 | mCherry | 28.9 / 26.2 |
| | H154 | Edamame | AAV9 | GFP | 29.3 / 31.9 |
| | | | CPP.16 | mCherry | 28.9 / 27.0 |
| | H155 | Okura | AAV9 | mCherry | 27.7 / 27.3 |
| | | | CPP.16 | GFP | 34.5 / 39.0 |
| | H188 | Umi | AAV9 | GFP | 32.2 / 33.9 |
| | | | BR1N | mCherry | 29.3 / 25.6 |
| $3.9 \times 10^{12}$ | H289 | Yoru | AAV9 | GFP | 25.3 / 26.1 |
| | | | CAP-Mac | mCherry | 30.0 / 27.7 |
| $5.0 \times 10^{12}$ | H057 | Kasumi | AAV9 | GFP | 23.2 / 22.7 |
| | I4471 | Ren | AAV9 | GFP | 16.2 / 19.0 |
| | H058 | Mitsuba | PHP.B | GFP | 22.4 / 26.2 |
| | I4466 | Suigetsu | PHP.B | GFP | 20.4 / 22.4 |
| $8.0 \times 10^{12}$ | H197 | Kuri | AAV9 | GFP | 47.9 / 50.1 |
| | | | CPP.16 | mCherry | 49.9 / 44.4 |
| $1.0 \times 10^{13}$ | H137 | Touga | PHP.eB | GFP | 18.7 / 25.1 |
| | H148 | Tougetsu | PHP.eB | GFP | 16.5 / 21.5 |
| | H377 | Bon | CAP-B10 | GFP | 19.4 / 21.1 |
| | H318 | Serori | AAV9 | GFP | 36.9 / 39.2 |
| | | | AAV-IR | mCherry | 32.1 / 29.4 |
| | H319 | Paseri | AAV9 | GFP | 38.4 / 42.0 |
| | | | AAV-Ly6H | mCherry | 44.8 / 43.1 |
| | SMP691 | Kinmokusei | AAV9 | mCherry | 33.2 / 32.1 |
| | | | VCAP-102 | GFP | 188.8 / 187.1 |
| | H365 | Nodoka | AAV9 | mCherry | 36.1 / 32.8 |
| | | | VCAP-102 | GFP | 229.4 / 213.2 |
| | H372 | Nana | VCAP-102 | GFP | 248.6 / 241.4 |
| $2.0 \times 10^{13}$ | H140 | Sekka | AAV9 | GFP | 21.6 / 22.6 |
| | H141 | Neyuki | PHP.eB | GFP | 35.9 / 41.3 |

Mean pixel intensity (0–255 a.u.) was quantified on the dorsal ventral surface of the whole brain approximately 4 weeks after intravenous delivery of each AAV vector. For animals receiving co-injections, fluorescence from AAV9 (GFP or mCherry) serves as the internal control for the paired variant. The table lists, for each individual: AAV dose (vg/kg), animal ID, name, capsid, expressed fluorophore, and mean dorsal and ventral fluorescence intensity. Measurements were performed under standardized exposure settings (GFP: 1 s; mCherry: 0.5 s).
